# Isolation and characterization of *Sporomusa carbonis* sp. nov., a carboxydotrophic hydrogenogen in the genus of *Sporomusa* isolated from a charcoal burning pile

**DOI:** 10.1101/2024.06.21.600036

**Authors:** Tim Böer, Florian P. Rosenbaum, Lena Eysell, Volker Müller, Rolf Daniel, Anja Poehlein

## Abstract

A Gram-negative bacterial strain, designated ACPt^T^, was isolated from the covering top soil of an active charcoal burning pile. Cells of ACPt^T^ were strictly anaerobic, rod-shaped and grew optimally at 40°C and pH 7. The substrates ribose, glucose, sucrose, raffinose, melezitose, pyruvate, vanillate, syringate, methanol and CO were utilized for growth. Phylogenomic analysis of the 4.1 Mb genome showed that strain ACPt represented a novel species of the genus *Sporomusa*. The most closely related species to ACPt^T^ was *Sporomusa malonica* with an average amino acid identity of 80.1%. The genome of ACPt^T^ encoded cytochromes, ubiquinones, the Wood-Ljungdahl gene cluster and an Rnf complex, which were identified as common features all *Sporomusa* type strains. However, strain ACPt did not ferment H_2_ + CO_2_ via acetogenesis as other *Sporomusa* species, but employed the metabolism of a carboxydotrophic hydrogenogen converting CO to H_2_ + CO_2_. Based on the genomic, morphological and physiological features presented in this study, strain ACPt^T^ is proposed as a novel species in the genus *Sporomusa* with the name *Sporomusa carbonis* sp. nov. (DSM 116159^T^ and CCOS 2105^T^).

## Introduction

The genus *Sporomusa* was formed in 1984 by the description of *Sporomusa ovata* isolated from sugar beet leaf silage and *Sporomusa sphaeroides* isolated from mud of the Leine river [1]. Hitherto, seven other *Sporomusa* species were validly published under the ICNP [2] comprising *Sporomusa paucivorans* [3], *Sporomusa termitida* [4], *Sporomusa acidovorans* [5], *Sporomusa malonica* [6], *Sporomusa silvacetica* [7], *Sporomusa aerivorans* [8] and *Sporomusa rhizae* [9]. Bacteria of the genus *Sporomusa* are morphologically characterized by their motile, curved rod-shaped and endospore-forming cells. Furthermore, *Sporomusa* cells stain Gram-negative and contain a multilayered cell wall [1]. Representatives of the genus *Sporomusa* are well known for utilizing H_2_ + CO_2_, alcohols (such as methanol, propanol and butanol), N-methylated compounds (such as sarcosine and betaine) and methoxylated monoaromates (such as vanillate and syringate) for growth [10]. H_2_ + CO_2_ is converted via acetogenesis using the Wood-Ljungdahl pathway (WLP), while the growth on alcohols, N-methylated and methoxylated compounds occurs by transferring methyl groups via a methyltransferase system and subsequent feeding of the methyl groups into the methyl-branch of the WLP [11]. Autotrophic growth on CO has only been reported for *S. ovata* [12] and *S. termitida* [4]. Energy conservation during acetogenesis in the mesophilic genus *Sporomusa* is achieved by building a chemiosmotic H^+^-gradient using an Rnf complex, in contrast to thermophilic acetogens utilizing an Ech complex [13]. The H^+^-gradient is subsequently utilized by an ATP synthase for the phosphorylation of ADP to ATP [14]. The genus *Sporomusa* comprises exclusively acetogenic and mesophilic strains containing both cytochromes b/c and ubiquinones. The presence of cytochromes and quinones is a rare trait of acetogens and besides *Sporomusa* members only present in acetogens of the thermophilic *Moorella* genus. Cytochromes and quinones were hypothesized to offer alternative routes for energy conservation during acetogenesis using the HdrABCMvhD or Fix complex [13,15]. Recently, *S. ovata* was shown to employ a three subunit *Sporomusa*-type Nfn transhydrogenase (Stn) instead of the two subunit Nfn transhydrogenase used by other acetogens for linking the redox pool of NADH and reduced ferredoxin during autotrophy to the cellular redox pool (NADPH) [16]. Biotechnological applications of bacteria from the genus *Sporomusa* have focused on the CO_2_-based production of bulk chemicals and biofuels via microbial electrosynthesis [17–22] or via a bacterial CdS biohybrid system [23]. *S. ovata* is particularly suitable for microbial electrosynthesis, as it exhibited some of the lowest H_2_-threshold found in described for acetogens [24]. Another biotechnological application is the transformation of cheap waste substrates such as methanol as an industrial feedstock for the production of biocommodities [25,26]. For the fermentation of CO-rich industrial waste gases, acetogens can be grown in co-culture with a carboxydotrophic hydrogenogenic bacterium initially converting CO with the water-gas shift reaction to H_2_ + CO_2_. Produced H_2_ + CO_2_ is subsequently fermented by an acetogen to acetate. For instance, the conversion of CO and acetate production of *S. ovata* could be significantly increased when grown in co-culture with *Citrobacter amalonaticus* [27]. The recent implementation of genetic tools for *S. ovata* have further paved the way for the utilization of *Sporomusa* members as industrial platform organisms [28]. All these applications of *Sporomusa* as a biocatalyst offer a renewable and sustainable route for the production of biocommodities, while simultaneously fixing the greenhouse gas CO_2_. Here, we present the description of the first carboxydotrophic hydrogenogen isolated from the genus *Sporomusa* and provide an analysis of high-quality genomes of *Sporomusa* type strains focused on genes and complexes potentially employed during lithotrophic growth.

## Methods

### Enrichment, isolation and cultivation

A sample from the top of the covering soil of an active burning charcoal pile in Hasselfelde, Germany (51°43’18.6’N 10°53’48.6’E) was taken in October 2021. The sample (1 g) was dissolved (50% w/v) in PBS-buffer (NaCl, 8 g/l; KCl, 0.2 g/l; Na_2_HPO_4_, 1.42 g/l; KH_2_PO_4_, 0.24 g/l) by shaking at 50 rpm for 1 h on a horizontal shaker (Adolf Kühner, Birsfelden, Swiss). Subsequently, the solution was pasteurized at 80°C for 10 min and 20 µl were used for the inoculation of enrichment cultures. The enrichment was performed in sterile anaerobic Hungate tubes filled with 10 ml of a modified DSM 311c minimal medium substituting sulfate salts with chloride salts, reducing yeast extract to 0.05 g/l and omitting fructose and peptone. Syringate (5 mM) was added to the media as a substrate prior to autoclaving. The enrichment culture was incubated at 35°C and samples were taken after 96 h and 264 h for cryoconservation (700 µl) by mixing the culture with 300 µl of glycerol (50%). Isolation was performed in the DSM 311c minimal medium from the cryo stock taken after 264 h by inoculating two subsequent serial dilutions of cultures in media supplemented with methanol (300 mM). The highest dilution showing growth was used to inoculate the subsequent dilution series. After isolation, the isolate was routinely cultivated in the modified DSM 311c medium of the enrichment cultures, but as a complex medium with yeast extract (2 g/l) and peptone (2 g/l). D-glucose (30 mM) was added as a substrate from a sterile anaerobic stock solution after media autoclaving. Reference *Sporomusa* strains were grown in the modified DSM 311c complex medium inoculated from lyophilised cultures obtained from the German Collection of Microorganisms and Cell Cultures (DSMZ, Braunschweig. Germany). *S. acidovorans*, *S. aerivorans*, *S. paucivorans*, *S. rhizae* were grown with methanol (300 mM), while *S. silvacetica* and *S. malonica* were grown with D-fructose (30 mM), both substrates were added from sterile anaerobic stock solutions after media autoclaving.

### Physiological and morphological characterization

Temperature, pH and NaCl optima were determined by cultivation in the modified DSM 311c complex medium supplemented with D-glucose (30 mM) inoculated to a start OD_600_ of 0.01. Growth was measured as optical density at 600 nm in Hungate tubes in triplicates after 48 h of incubation using the spectrophotometer WPA S800 (Biochrom, Berlin, Germany). The temperature optimum was investigated by incubation at temperatures ranging from 10 to 55°C in 5°C intervals. NaCl and pH optima were investigated at the respective temperature optima of the strains. NaCl tolerance was investigated with added NaCl concentrations of 0%, 0.1%, 1%, 2%, 3%, 4% and 5%. The pH optimum was investigated by setting the pH to 5.5 to 9 in 0.5 intervals using HCl/NaOH. Gram staining was carried out with the Claus protocol [29]. Substrate utilization was evaluated by cultivation in Hungate tubes with the modified DSM 311c minimal medium adding substrates from sterile anaerobic stock solutions in a final concentration of 30 mM. Vanillate (5 mM) and syringate (5 mM) were added to the media before autoclaving. Growth was measured from triplicate cultures after 48 h of incubation at the temperature optimum of the respective strain. Substrates not being used in the initial 48 h of incubation were continuously assessed for a total of 2 weeks before substrate utilization was considered negative. The cellular fatty acid profile was determined by the identification service provided by the DSMZ. Cell morphology was investigated with a Jeol 1011 transmission electron microscope (Georgia Electron Microscopy, Freising, Germany) and negatively stained cells. Negative cell staining was achieved by mixing 5 µl cell suspension with 5 µl phospho-tungstic acid solution (0.5% w/v). The mixture was transferred to a vaporized carbon mica and subsequently placed on a copper-coated grid (PLANO GmbH, Marburg, Germany). Concentrations of acetate, CO and H_2_ were determined as described in Weghoff et al. 2016 [30,31].

### Genome sequencing, assembly and analysis

DNA isolation and subsequent Illumina sequencing were performed as described in Böer et al. [32]. For Nanopore sequencing high-molecular weight DNA was isolated from a separately cultivated batch with the Monarch Genomic DNA Purification kit (New England Biolabs, Frankfurt, Germany) and library preparation was performed with 1.5 µg using the ligation sequencing kit 1D (SQK-NBD114.24) and the native barcode expansion kit (EXP-NBD104) as recommended by the manufacturer (Oxford Nanopore Technologies, Oxford, UK). Nanopore sequencing was conducted for 72 h employing the MinION device Mk1B, the SpotON flow cell R10.4.1 and the MinKNOW software (v23.4.6) following the instructions of the manufacturer (Oxford Nanopore Technologies). Demultiplexing and base calling of Nanopore sequencing data were performed with the Dorado (v6.5.7) software in SUP mode. Long-read (Nanopore reads) de novo genome assemblies were performed with trycycler as outlined by Wick et al. [33]. Quality control and adapter trimming of paired-end Illumina sequences was performed with Fastp (v0.23.4) [34] and Trimmomatic (v0.39; LEADING: 3, TRAILING: 3, SLIDINGWINDOW:4:15, MINLEN:50) [35]. Adapter trimming of Nanopore sequencing data was performed with Porechop (v0.2.4) and sequences were subsequently quality filtered with Filtlong (v0.2.1) to a minimal read length of 1 kb and 5% of the sequences with the worst quality score were discarded. Filtered Nanopore reads were subsampled 24 times with Trycycler (v0.5.4) [33] and eight subsets each were used as input for the assemblers Flye (v2.9.2) [36], Canu (v2.2) [37] and Raven (v1.8.3) [38]. Assemblies were combined to a single consensus sequence with Trycycler and the consensus sequence was polished with the full Nanopore data using Medaka (v1.10.0) and finally polished with the processed short-read sequences using Polypolish (v0.5.0) [39]. Genomes were reorientated manually to start with the *dnaA* gene. Genome annotations were performed with Prokka (v1.14.6) [40] and quality assessment of the final genome assemblies was conducted with CheckM2 (v1.0.2) [41]. Visualization of gene clusters were performed with clinker (v.0.0.28) [42], genes showing a protein sequence identity of ≥50% were assigned the same color. Phylogenomic analysis was carried out by determining the average nucleotide identity (ANI) using orthoANI (v0.5.0) [43], digital DNA-DNA hybridization values (dDDH) of the formula d_4_ using the Genome to Genome Distance Calculator (v3.0) [44], the percentage of conserved proteins (POCP) using the POCP pipeline (v2.3.2) [45] based on the method described by Quin et al. [46] and the average amino acid identity (AAI) using ezAAI (v1.2.3) [47]. The whole genome sequence-based phylogram was obtained by using the Type (Strain) Genome Server (TYGS) [48]. The core/pan genome of the *Sporomusa* genus and strain specific unique orthologous groups (OGs) were determined with Proteinortho (v6.3.0) [49] applying an identity threshold of ≥50% and an e-value of ≤1e^-10^. Unique OGs were visualized in R with the ggplot2 package (v3.4.1) [50]. The 16S rRNA gene sequence was amplified with a DreamTaq polymerase using genomic DNA as the template and the primers 08F (5′-AGAGTTTGATCCTGGC-3′) and 1504R (5′-TACCTTGTTACGACTT-3′) following the standard instructions of the manufacturer (Thermo Fisher Scientific, Waltham, MA, USA). The PCR product was purified using the Master Pure Complete DNA & RNA Purification Kit (Epicentre, Madison, USA) and Sanger sequenced by Seqlab (Microsynth Seqlab GmbH, Göttingen, Germany).

## Results and Discussion

### Genomic and phylogenomic analysis

The genome assembly and annotation data for the *Sporomusa* genomes is summarized in Table 1. Genome sizes ranged from 4.1 to 6.5 Mb and average GC-content from 43 to 49.1%. With 4.1 Mb the genome size of isolate ACPt was considerably smaller than the reference *Sporomusa* genomes (4.8 to 6.5 Mb) and the *Methylomusa anaerophila* genome (4.8 Mb). With an average GC-content of 46.2%, the genome of strain ACPt fell within the average GC-content of other *Sporomusa* genomes ranging from 43.0 to 49.1% and was similar to the average GC-content of the *M. anaerophila* genome with 46.6%. Correlating with the genome size, the lowest number of putative genes was encoded by the genome of the ACPt isolate (4,148), while the highest number of genes was encoded by *S. aerivorans* (6,303).

**Table 1.** Genome and annotation data of the novel ACPt strain, the *Sporomusa* type strains of *S. malonica*, *S. aerivorans*, *S. termitida*, *S. paucivorans*, *S. rhizae*, *S. sphaeroides*, *S. ovata*, *S. acidovorans*, *S. silvacetica* and the type strain *M. anaerophila*.

|  | ACpt * | *S. malonica* * | *S. aerivorans* * | *S. termitida* ** | *S. paucivorans* * | *S. rhizae* * | *S. sphaeroides* *** | *S. ovata* *** | *S. acidovorans* * | *S. silvacetica* * | *M. anaerophila* **** |
| --- | --- | --- | --- | --- | --- | --- | --- | --- | --- | --- | --- |
| Culture collection identifier | DSM 116159<sup>T</sup><br>CCOS 2105<sup>T</sup> | DSM 5090<sup>T</sup> | DSM 13326<sup>T</sup> | DSM 4440<sup>T</sup> | DSM 3697<sup>T</sup> | DSM 16652<sup>T</sup> | DSM 2875<sup>T</sup> | DSM 2662<sup>T</sup> | DSM 3132<sup>T</sup> | DSM 10669<sup>T</sup> | JCM 31821<sup>T</sup> |
| Genome Size (bp) | 4,144,804 | 5,312,210 | 6,496,238 | 5,184,890 | 4,787,310 | 5,829,736 | 4,956,256 | 5,433,971 | 5,955,502 | 6,046,356 | 4,781,198 |
| Plasmid Size (bp) | - | - | 15,390- | 131,118 | - | - | 59,087 | - | - | - | - |
| GC-content (%) | 46.2 | 44.6 | 46.7 | 49.1 | 47.3 | 43.3 | 47.2 | 43.3 | 44.7 | 43.0 | 46.6 |
| CheckM2 completeness/contamination scores (%) | 100/2.90 | 100/5.91 | 100/12.26 | - | 99.97/1.53 | 100/9.66 | 99.99/1.19 | 100/7.57 | 100/6.45 | 100/6.41 | - |
| Genes | 4,148 | 5,118 | 6,303 | 4,872 | 4,483 | 5,495 | 4,656 | 5,314 | 5,750 | 5,913 | 4,268 |
| CDS | 4,022 | 4,955 | 6,136 | 4,735 | 4,346 | 5,324 | 4,511 | 5,112 | 5,604 | 5,757 | 4,183 |
| Functional proteins | 2,171 | 2,757 | 3,255 | 3,659 | 2,551 | 2,965 | 2,596 | 3,609 | 3,056 | 3,015 | 3,083 |
| Hypothetical proteins | 1,851 | 2,198 | 2,881 | 1,076 | 1,795 | 2,359 | 1,915 | 1,503 | 2,548 | 2,742 | 1,100 |
| rRNA (5S; 16S; 23S) | 5, 11, 10 | 18, 16, 14 | 10, 14, 12 | 9, 12, 10 | 10, 11, 10 | 13, 14, 12 | 10, 12, 11 | 15, 15, 12 | 9, 11, 10 | 13, 14, 12 | 4, 4, 3 |
| tRNAs | 99 | 114 | 130 | 105 | 105 | 131 | 111 | 160 | 115 | 116 | 73 |
| tmRNAs | 1 | 1 | 1 | 1 | 1 | 1 | 1 | 1 | 1 | 1 | 1 |
| CRISPR repeats | 4 | 0 | 0 | 9 | 3 | 5 | 2 | 3 | 6 | 2 | 9 |
| BioProject genome | PRJNA1082235 | PRJNA1081642 | PRJNA1081645 | - | PRJNA1081640 | PRJNA1081641 | - | - | PRJNA310710 | PRJNA310713 | - |
| BioSample genome | SAMN40201554 | SAMN40183415 | SAMN40183414 | - | SAMN40182996 | SAMN40183110 | - | - | SAMN04455122 | SAMN04455153 | - |
| Sequence Read Archive Illumina genome | SRR28210317 | SRR28149488 | SRR28161548 | - | SRR28148131 | SRR28148136 | - | - | SRR28147534 | SRR28762162 | - |
| Sequence Read Archive Nanopore genome | SRR28210318 | SRR28161377 | SRR28161549 | - | SRR28148125 | SRR28149486 | - | - | SRR28148124 | SRR28762299 | - |
| GenBank assembly accession |  |  |  | GCA_007641255.1 |  |  | GCA_001941975.2 | GCA_000445445.2 |  |  | GCA_003966895.1 |
| GenBank 16S rRNA | PP693464 | - | - | - | - | - | - | - | - | - | - |

\* this study \*\* [51] \*\*\* Böer et al. 2024 \*\*\*\* [52]

Data for the phylogenomic classification of the ACPt isolate to type strains of the genus *Sporomusa* and *Methylomusa* are summarized in Table 2. The phylogenomic comparison of strain ACPt with *Sporomusa* type strains and *M. anaerophila* showed orthoANI values ranging from 70.9% to 76.3% and dDDH (d_4_) [53] values ranging from 20.5% to 21.4%. As these values fell below the species threshold of under 95% for ANI [43,54] and under 70% for dDDH, the ACPt isolate represented a novel species in the genus *Sporomusa*. In order to determine whether the ACPt isolate could represent a novel genus in the family of *Sporomusaceae*, we additionally calculated POCP and AAI values. POCP values ranged from 56.4% to 63.3% and AAI values from 71.3% to 80.1%. POCP values of under 50% [55] and AAI values of 76% [56] were recommended as thresholds for genus delineation. In all AAI comparisons of the ACPt genome to *Sporomusa* reference genomes, we obtained values directly below and over the recommended genus threshold of 76%. The AAI comparison to the genome of *M. anaerophila* was with 71.3% considerably lower than the threshold of 76%. This indicated that strain ACPt should be classified as a novel species of the genus *Sporomusa*. POCP comparisons of genomes from the genus *Sporomusa* yielded values considerably higher than the recommended 50% threshold for genus delineation. However, with a POCP value of 56.4% this also applied for the comparison of strain ACPt with the *M. anaerophila* genome. This indicated that the 50% POCP threshold for genus delineation was not applicable for this phylogenomic cluster. The most closely related *Sporomusa* species was *S. malonica* with a POCP value of 63.3% and an AAI value of 80.1%. Correspondingly, based on the complete phylogenomic analysis strain ACPt was classified as a novel species in the genus *Sporomusa* and the name *Sporomusa carbonis* is proposed. The whole genome-based phylogram placed the genome of strain ACPt into a phylogenomic group with the type strains of *S. malonica* and *S. termitida* (Figure 1).

**Table 2.** Phylogenomic analysis of strain ACPt, the *Sporomusa* type strains of *S. malonica*, *S. aerivorans*, *S. termitida*, *S. paucivorans*, *S. rhizae*, *S. sphaeroides*, *S. ovata*, *S. acidovorans*, *S. silvacetica* and the type strain *M. anaerophila*.

|  | orthoANI (%) | dDDH (d<sub>4</sub>) | POCP (%) | AAI (%) |
| --- | --- | --- | --- | --- |
| *S. malonica* | 76.3 | 21.4 | 63.3 | 80.1 |
| *S. aerivorans* | 74.2 | 20.7 | 58.0 | 77.3 |
| *S. termitida* | 74.3 | 20.9 | 61.8 | 77.2 |
| *S. paucivorans* | 74.7 | 21.2 | 63.5 | 76.8 |
| *S. rhizae* | 73.8 | 20.5 | 58.7 | 76.7 |
| *S. sphaeroides* | 74.9 | 21.0 | 61.8 | 76.5 |
| *S. ovata* | 74.0 | 21.2 | 59.1 | 75.9 |
| *S. acidovorans* | 73.5 | 20.2 | 61.0 | 75.8 |
| *S. silvacetica* | 73.9 | 21.2 | 57.4 | 75.7 |
| *M. anaerophila* | 70.9 | 21.1 | 56.4 | 71.3 |

**Fig. 1.**
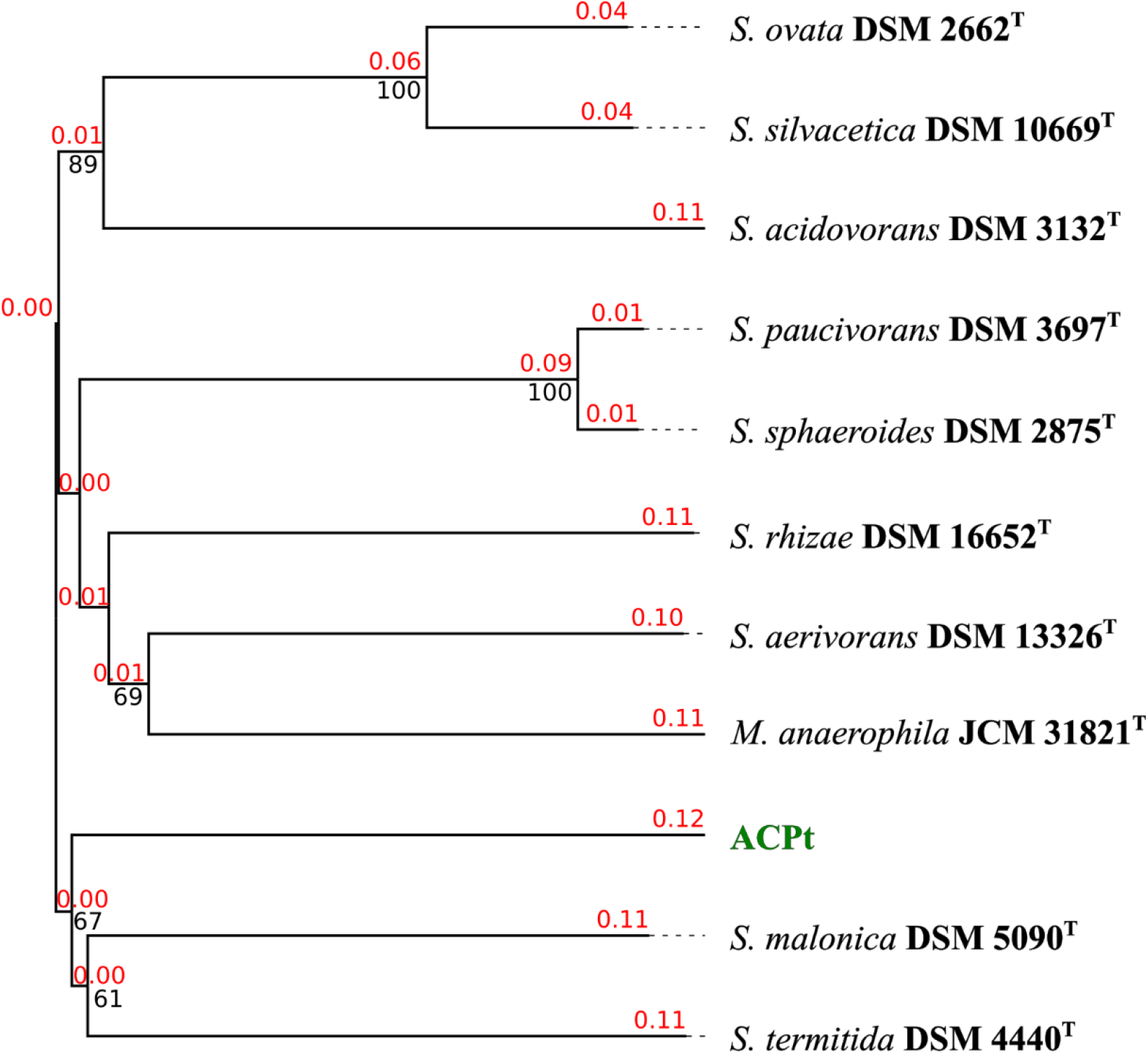
Whole genome sequence-based phylogram of the novel ACPt isolate and the type strains of the genera *Sporomusa* and *Methylomusa*. Branch lengths are scaled in terms of GBDP distance formula *d_5_*. Red numbers above branches are GBDP pseudo-bootstrap support values > 60% from 100 replications, with an average branch support of 71.5%. Black numbers below branches are confidence scores. The tree was rooted at the midpoint.

Core/pan genome analysis of the genus *Sporomusa* identified a pan genome of 16,103 OGs and a core genome of 1,630 OGs. The highest amount of unique OGs were identified for *S. silvacetica* (1,190), *S. aerivorans* (1,182), *S. acidovorans* (1,172), *S. rhizae* (1,099) and *S.ovata* (1,061), while lower amount of unique OGs were identified for strain ACPt (857), *S. termitida* (741), *S. malonica* (705), *S. sphaeroides* (431) and *S. paucivorans* (244) (Figure 2). Unique OGs identified in the genome of strain ACPt comprised for instance a long-chain primary alcohol dehydrogenase (AdhA, SCACP_29740), a ferredoxin-NADP reductase (Fpr, SCACP_12760), a ferredoxin (Fer, SCACP_12770), a bifunctional homocysteine S-methyltransferase/5,10-methylenetetrahydrofolate reductase (YitJ, SCACP_12780), a pyruvate carboxylase (PycA, SCACP_22850) and a gene cluster encoding subunits of a CO-oxidizing/H_2_-evolving enzyme complex (CooMKLXUHFSC, SCACP_37230-37350). The latter gene cluster was most similar to the gene clusters of carboxydotrophic hydrogenogens (Ech group 4c according to the classification by Schoelmerich et al. 2020. [57]) from the thermophilic genera *Thermincola*, *Thermoanaerosceptrum, Thermosinus, Carboxydocella, Carboxydothermus* and *Calderihabitans* (Figure 3).

**Fig. 2.**
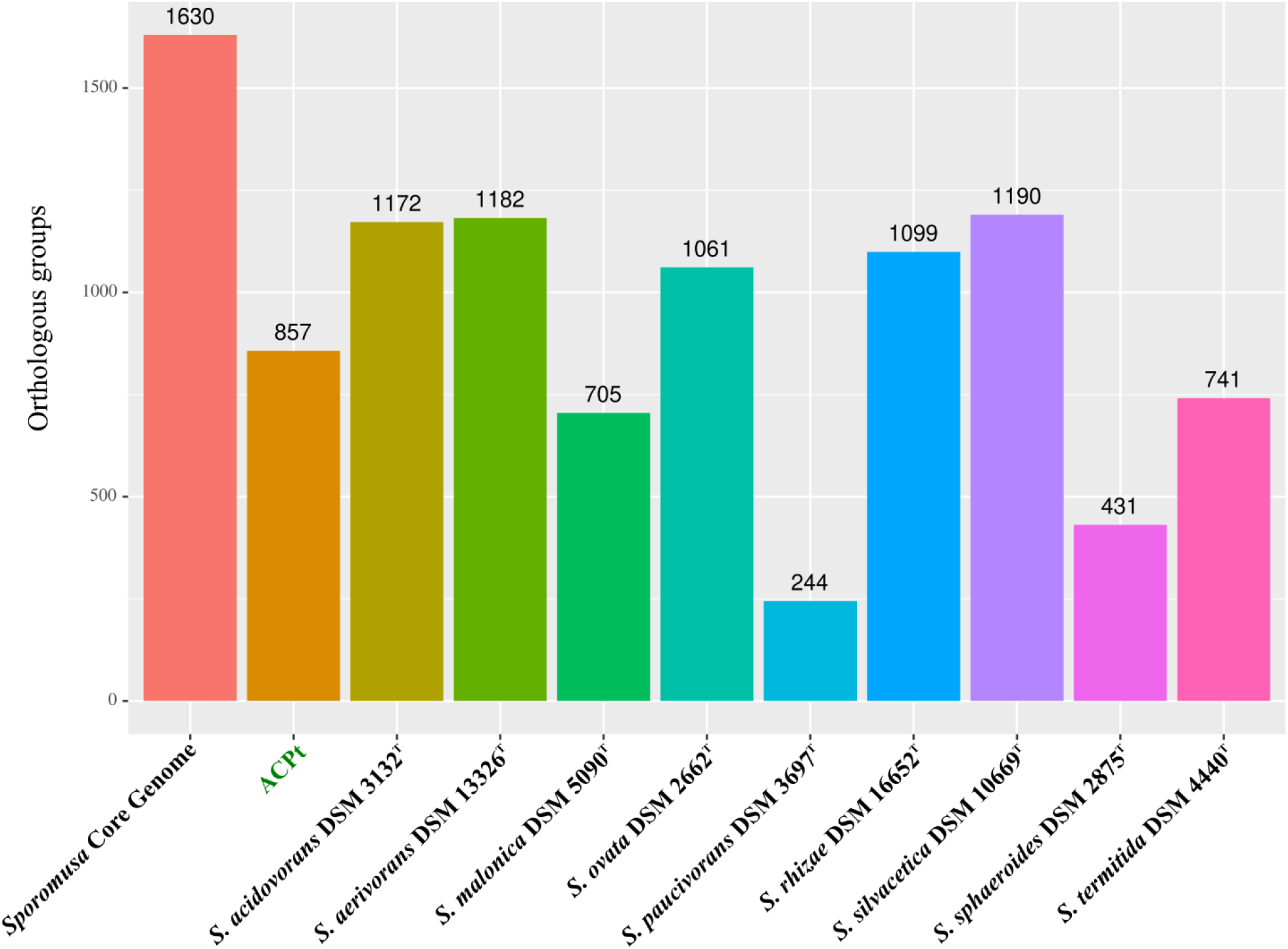
Pan/core genome analysis of the genus *Sporomusa* including the genomes of all *Sporomusa* type strains.

**Fig. 3.**
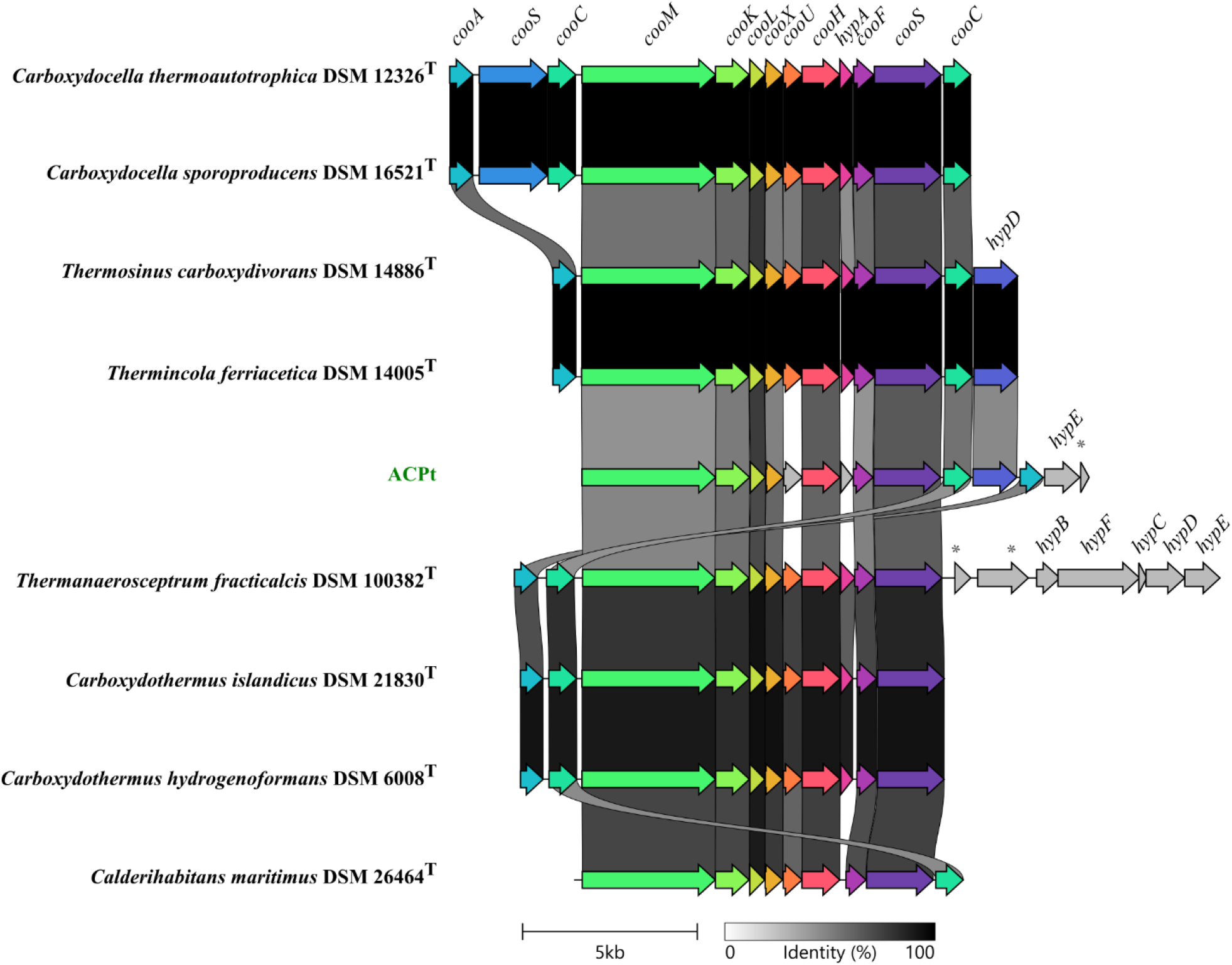
Structure of the gene cluster of the monofunctional carbon monoxide dehydrogenase in strain ACPt in comparison to carboxydotrophic hydrogenogens of thermophiles. Genes encoding proteins with 50% sequence identity or higher were given the same color. The following gene abbreviations were used: *cooA*, transcriptional regulator CooA; *cooS*, monofunctional carbon monoxide dehydrogenase CooS; *cooC*, carbon monoxide dehydrogenase accessory protein; *cooM*, carbon monoxide dehydrogenase complex subunit CooM; cooK, carbon monoxide dehydrogenase complex subunit CooK; *cooL*, carbon monoxide dehydrogenase complex subunit CooL; *cooX*, carbon monoxide dehydrogenase complex subunit CooX; cooU, carbon monoxide dehydrogenase complex subunit CooU; *cooH*, carbon monoxide dehydrogenase complex subunit CooH; hypA, hydrogenase maturation factor HypA; *cooF*, ferredoxin type subunit of carbon monoxide dehydrogenase complex; *hypD*, hydrogenase maturation factor HypD; *hypE*, carbamoyl dehydratase HypE; *, hypothetical protein; hypB, hydrogenase maturation factor HypB; *hypF*, carbamoyltransferase HypF; *hypC*, hydrogenase maturation factor HypC

All *Sporomusa* genomes encoded an Rnf complex, the WLP-gene cluster, a monofunctional carbon monoxide dehydrogenase (CooS), cytochromes b and c, ubiquinones, an HdrABC-mvhD complex (Hdr) and a FixABCX complex (Fix). The *Sporomusa* type Nfn transhydrogenase (Stn) was encoded by the genomes of *S. ovata*, *S. silvacetica* and *M. anaerophila*, while other *Sporomusa* species harbored instead genes coding for a three subunit electron-bifurcating hydrogenase (HydABC) at this genome location. *Sporomusa* species lacking the Stn transhydrogenase contained genes for the Nfn transhydrogenase [16,58]. An overview of the detected genes and complexes potentially involved in lithotrophic growth of the genus *Sporomusa* is shown in Table 4.

The structure of the genomic region encoding the WLP gene cluster and adjacent genes in the genus *Sporomusa* and *Methylomusa* are visualized in Figure 4. The structure of the WLP gene cluster including the genes encoding the Hdr complex were highly conserved in *Sporomusa* and *Methylomusa* members. The genome of strain ACPt additionally encoded a ferredoxin-NADP reductase gene (*fpr*) in synteny with a ferredoxin (*fer*) and a bifunctional homocysteine S-methyltransferase/5,10-methylenetetrahydrofolate reductase (*yitJ*) upstream of the WLP gene cluster. The structure of the genes downstream of the Hdr complex in the *Sporomusa* genomes differed by the presence of the HydABC hydrogenase or the StnABC transhydrogenase. They also differed in the type and amount of formate dehydrogenases being encoded. All *Sporomusa* genomes coded for additional hydrogenase subunit genes (*hydA, stnB, hndB)* in synteny with several genes for hypothetical proteins at the end of this genomic region.

**Fig. 4.**
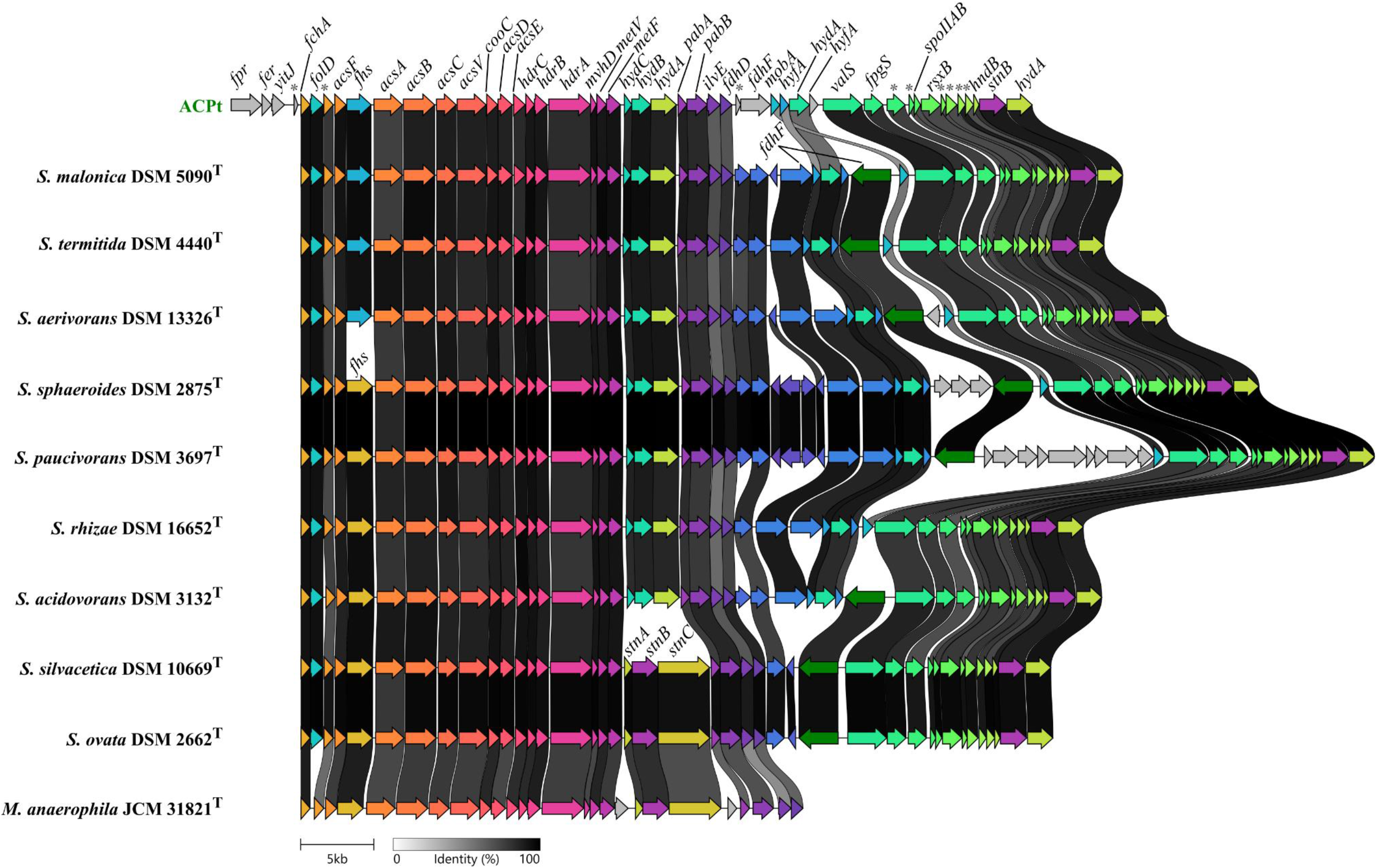
Structure of the Wood-Ljungdahl gene cluster in type strains of the genus *Sporomusa* and *Methylomusa*. Genes encoding proteins with 50% sequence identity or higher were given the same color. The following gene abbreviations were used: *fpr*, ferredoxin-NADP reductase; *fer*, ferredoxin ;*yitJ*, bifunctional homocysteine S-methyltransferase/5,10-methylenetetrahydrofolate reductase; *\**, hypothetical protein; *fchA*, methenyl THF cyclohydrolase; *folD*, bifunctional cyclohydrolase/dehydrogenase; *acsF*, carbon monoxide dehydrogenase accessory protein; *fhs*, formyl THF synthetase; *acsA*, anaerobic carbon-monoxide dehydrogenase catalytic subunit; *acsB*, carbon monoxide dehydrogenase/acetyl-CoA synthase subunit beta; *acsC*, CoFeSP large subunit; *acsV*, corrinoid activation/regeneration protein; *cooC*, carbon monoxide dehydrogenase accessory protein; *acsD*, CoFeSP small subunit; *acsE*, methyl THF CoFeSP methyltransferase; *hdrC*, heterodisulfide oxidoreductase iron-sulfur cluster-binding subunit; *hdrB*, heterodisulfide reductase subunit B; *hdrA*, heterodisulfide reductase subunit A; *mvhD*, methyl-viologen-reducing hydrogenase delta subunit; *metV*, methylene THF reductase C-terminal catalytic subunit; *metF*, methylene THF reductase large subunit; *hydC*, electron bifurcating hydrogenase subunit HydC; *hydB*, electron bifurcating hydrogenase subunit HydB; *hydA*, electron bifurcating hydrogenase subunit HydA; *stnA*, *Sporomusa*-type Nfn transhydrogenase subunit A; *stnB*, *Sporomusa*-type Nfn transhydrogenase subunit B; *stnC*, *Sporomusa*-type Nfn transhydrogenase subunit C; *pabA*, aminodeoxychorismate/anthranilate synthase component 2; *pabB*, aminodeoxychorismate synthase component 1; *ilvE*, branched-chain-amino-acid aminotransferase; *fdhD*, sulfur carrier protein; *fdhF*, formate dehydrogenase H; *mobA*, molybdenum cofactor guanylyltransferase; *hyfA*, hydrogenase-4 component A; *valS*, valine--tRNA ligase; *fpgS*, folylpolyglutamate synthase; *spoIIAB*, anti-sigma F factor; *rsxB*, ion-translocating oxidoreductase complex subunit B; *hndB*, NADP-reducing hydrogenase subunit HndB.

### Morphological, physiological and chemotaxonomic characterization

An overview of the morphological and physiological characterization of strain ACPt and the reference data of other *Sporomusa* type strains is shown in Table 3. Cells of strain ACPt showed the *Sporomusa* characteristic curvature of the rod-shaped cells (Figure 5), stained Gram-negative and showed growth only under strict anaerobic conditions. Growth occurred in the temperature range of 35-50°C and was optimal at 40°C under the tested conditions. In comparison to the reported data for other *Sporomusa* species growing between 15 and 45°C, the temperature range of strain ACPt was shifted towards higher temperatures. With growth occurring at 50°C, strain ACPt showed the highest growth temperature reported for a *Sporomusa* type strain. Furthermore, strain ACPt grew optimally when NaCl was not added to the medium and at pH 7. The NaCl tolerance of other *Sporomusa* species has not been reported, the determined pH optimum of 7 was in line with the data of other *Sporomusa* species descriptions. Cellular fatty acid analysis identified Iso-βOH-C_13:0_ (15.6%), Iso-C_15:0_ (11.4%) and Iso-C_11:0_ (9.0%) as the predominant cellular fatty acids of strain ACPt. In comparison to the reported cellular fatty acid profile of *S. ovata* An4, *S. ovata* DSM 2662^T^ and *S. aerivorans* DSM 13326^T^ [12], strain ACPt contained higher amounts of C_15:0_, Iso-C_15:0_ and lower amounts of C_16:1_ Δ7 (Supplementary material, Table S1). Strain ACPt utilized ribose, fructose, glucose, sucrose, lactose, melezitose, pyruvate, vanillate, syringate and methanol for growth. Growth with glucose, sucrose, lactose and melezitose were not reported for any other *Sporomusa* species and were thereby identified as potential unique metabolic feature of strain ACPt. Arabinose, xylose, galactose, mannose, rhamnose, glucuronic acid, glycerol, sorbitol, mannitol, inositol, cellobiose, trehalose, maltose, melibiose, raffinose, formate, lactate, malate, 3-OH-butyrate, citrate, sarcosine, DMG, betaine, ethanol, propanol, 1,2-propanediol and butanol were not utilized for growth under the tested conditions. We did not observe the transformation of H_2_ + CO_2_ to acetate by strain ACPt under the tested conditions. Instead, strain ACPt transformed CO to the final product of H_2_ + CO_2_. These observations imply that strain ACPt employs the metabolism of a carboxydotrophic hydrogenogen transforming CO with the water-gas shift reaction to H_2_ + CO_2_, which is coupled to energy conservation of the produced ferredoxin at a membrane-associated Ech complex [59]. This hypothesis was further supported by the identification of the strain ACPt unique *CooMKLXUHFSC* gene cluster during the pan/core genome analysis of the genus *Sporomusa*. The metabolism of carboxydotrophic hydrogenogens was hypothesized to detoxify CO-exposed microenvironments [60]. In the charcoal burning pile environment strain ACPt was exposed to high concentrations of CO, which putatively led to the adaptation of an acetogenic bacterium to a carboxydotrophic hydrogenogen for the advantage of rapid detoxification. The *CooMKLXUHFSC* gene cluster was hypothesized to be propagated by horizontal gene transfer, as this genomic region shows high similarities between isolated strains of phylogenetically diverse taxa [60].

**Table 3.**
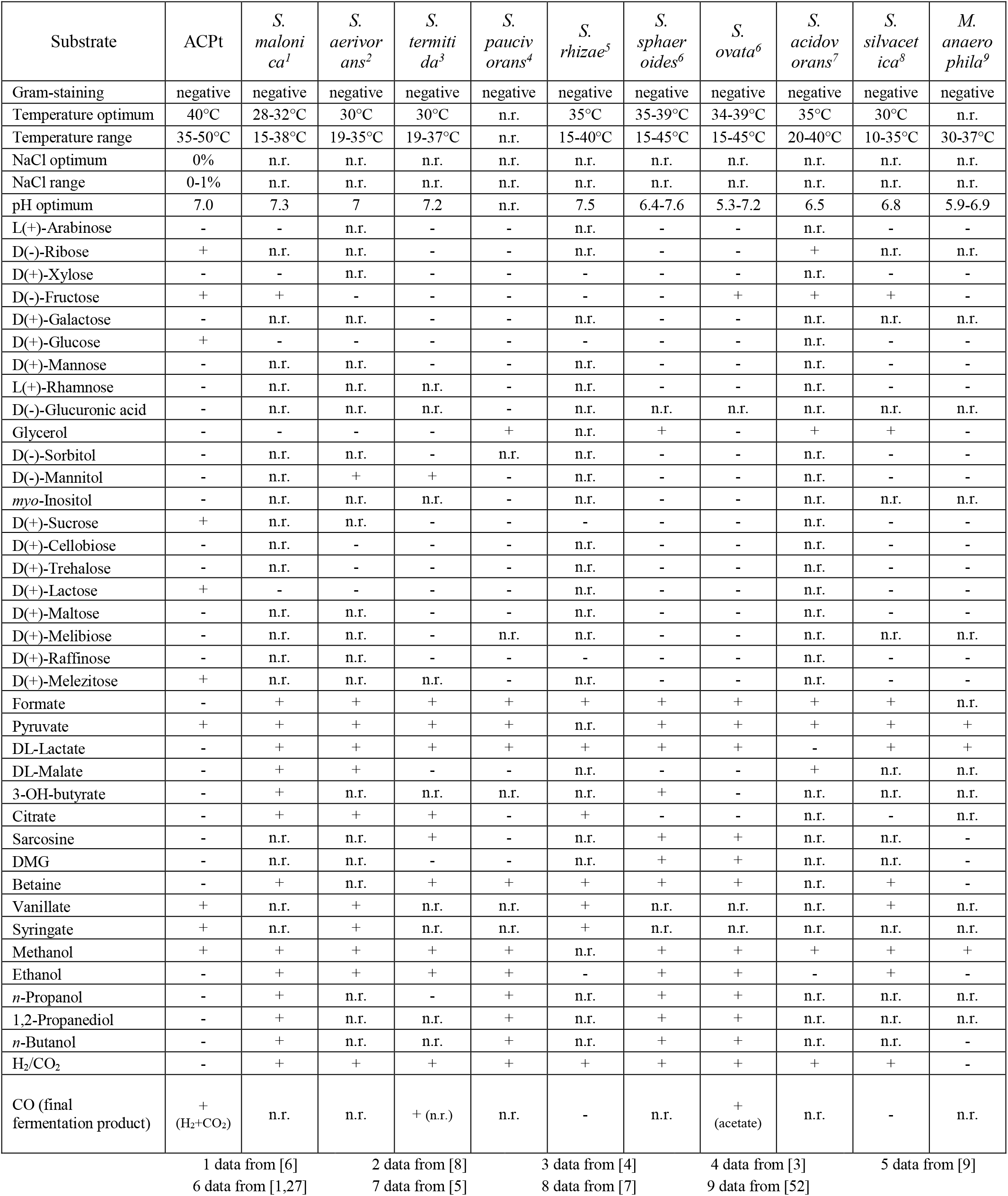
Morphological and physiological characterization of strain ACPt in comparison to the data reported for other *Sporomusa* type strains. +, positive; -, negative; n.r., not reported.

**Fig. 5.**
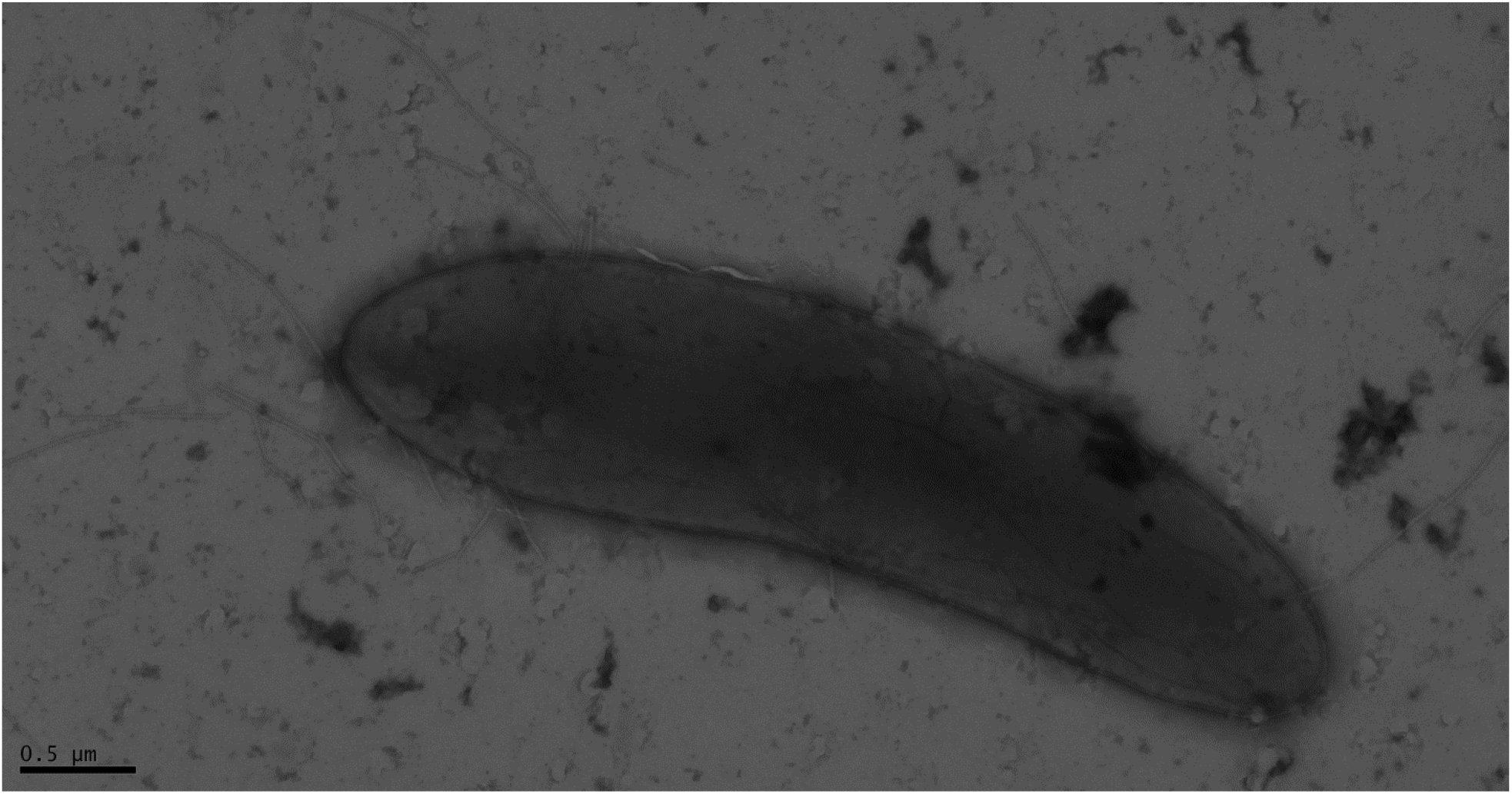
Transmission electron micrographs of negatively stained cells of strain ACPt.

**Table 4.** Genomic analysis of the type strains of the genera *Sporomusa* and *Methylomusa* with regards to the presence of the Rnf complex (Rnf), Ech complex (Ech), Wood-Ljungdahl pathway (WLP), monofunctional carbon monoxide dehydrogenase (CooS), cytochromes (Cyt), ubiquinones (UQ), HdrABC-mvhD complex (Hdr), FixABCX complex (Fix), NfnAB transhydrogenase (Nfn) and StnABC transhydrogenase (Stn). +, genes present; - genes absent.

|  | Rnf | Ech | WLP | CooS | Cyt b/c | UQ | Hdr | Fix | Nfn | Stn |
| --- | --- | --- | --- | --- | --- | --- | --- | --- | --- | --- |
| ACPt | + | + | + | + | + | + | + | + | + | - |
| *S. malonica* | + | - | + | + | + | + | + | + | + | - |
| *S. aerivorans* | + | - | + | + | + | + | + | + | + | - |
| *S. termitida* | + | - | + | + | + | + | + | + | + | - |
| *S. paucivorans* | + | - | + | + | + | + | + | + | + | - |
| *S. rhizae* | + | - | + | + | + | + | + | + | + | - |
| *S. sphaeroides* | + | - | + | + | + | + | + | + | + | - |
| *S. ovata* | + | * | + | + | + | + | + | + | - | + |
| *S. acidovorans* | + | - | + | + | + | + | + | + | + | - |
| *S. silvacetica* | + | - | + | + | + | + | + | + | - | + |
| *M. anaerophila* | + | - | + | + | + | + | + | - | - | + |

\* Ech-like complex, not in synteny of *cooS*

### Description of *Sporomusa carbonis* sp. nov

*Sporomusa carbonis* (car.bo’nis. L. gen. masc. n. *carbonis*, of coal, of charcoal). Cells show the form of a curved rod and stain Gram-negative. Growth occurs only under strict anaerobic conditions and ranges from 35-50°C (optimum 40°C) and added NaCl concentrations from 0-1% and optimal without addition. The optimal pH for growth was 7 and maximal growth occurs with glucose as a substrate. Furthermore, the substrates ribose, fructose, glucose, sucrose, lactose, melezitose, pyruvate, vanillate, syringate and methanol are utilized. CO is transformed to H_2_ and CO_2_ as the major fermentation product. Does not grow on H_2_ + CO_2_, arabinose, xylose, galactose, mannose, rhamnose, glucuronic acid, glycerol, sorbitol, mannitol, inositol, cellobiose, trehalose, maltose, melibiose, formate, lactate, malate, 3-OH-butyrate, citrate, sarcosine, DMG, betaine, ethanol, propanol, 1,2-propanediol and butanol. The predominant cellular fatty acids are Iso-βOH-C_13:0_ (15.6%), Iso-C_15:0_ (11.4%) and Iso-C_11:0_ (9.0%). The type strain genome comprises 4.1 Mbp and a GC-content of 46.2%. The type strain ACPt was isolated from the top of the covering soil of an active charcoal burning pile, Hasselfelde, Germany. The GenBank/EMBL/DDBJ accession number of the 16S rRNA gene is PP693464. The type strain of *S. carbonis* is ACPt^T^ and was deposited under the identifiers DSM 116159^T^ and CCOS 2105^T^.

## Author contributions

TB: Conceptualization, Data curation, Formal analysis, Investigation, Methodology, Visualization, Writing – original draft, Writing – review & editing

FR: Data Curation, Investigation, Validation, Writing – review & editing

LE: Investigation, Validation, Writing – review & editing

VM: Funding acquisition, Supervision, Validation, Writing – review & editing

RD: Conceptualization, Funding acquisition, Project administration, Supervision, Validation, Writing – review & editing

AP: Conceptualization, Funding acquisition, Methodology, Supervision, Validation, Writing – review & editing

## Acknowledgements

We thank Michael Hoppert, Ines Friedrich and Vanessa Beck for providing the transmission electron micrograph. Furthermore, we thank Melanie Heinemann and Mechthild Bömeke for technical assistance. We thank the Open Access Publication Funds of the University of Göttingen and the “Ministerium für Wissenschaft und Kultur” of Lower Saxony (Germany) for supporting publishing open access.

