## supplementary table S1 for "Isolation and characterization of *Sporomusa carbonis* sp. nov., a carboxydotrophic hydrogenogen in the genus of *Sporomusa* isolated from a charcoal burning pile"

### Supplementary Material

**Table S1.** Cellular fatty acid composition of *S. carbonis* in comparison to the reported data (Balk et al. 2010) for *S. aerivorans* DSM 13326$^{T}$, *S. ovata* DSM 2662$^{T}$ and *S. ovata* DSM 21435.

| Fatty acid | *S. carbonis* DSM 116159$^{T}$ | *S. aerivorans* DSM 13326$^{T*}$ | *S. ovata* DSM 2662$^{T*}$ | *S. ovata* DSM 21435$^{*}$ |
|---|---|---|---|---|
| $C_{11:0}$ | 0.5 | - | - | - |
| Iso-$C_{11:0}$ | 9.0 | 1.1 | 1.2 | 2.3 |
| Iso-$C_{11:0}$ DMA | 0.2 | - | - | - |
| Iso-βOH-$C_{11:0}$ | 0.3 | - | - | - |
| βOH-$C_{11:0}$ | - | 0.9 | 1.3 | - |
| βOH-$C_{12:0}$ | 1.4 | 10.5 | 7.3 | 1.5 |
| $C_{13:0}$ | 0.3 | - | - | - |
| $C_{13:1}$ Δ3 | 0.9 | - | - | - |
| Iso-$C_{13:0}$ | 0.2 | - | - | - |
| Iso-βOH-$C_{13:0}$ | 15.6 | 8.6 | 8.0 | 26.5 |
| βOH-$C_{13:0}$ | - | 2.0 | 4.1 | - |
| βOH-$C_{13:1}$ | - | 0.8 | 1.1 | - |
| $C_{14:0}$ | 3.0 | 1.1 | 1.2 | 0.5 |
| $C_{14:0}$ DMA/βOH-$C_{13:0}$/Iso-$C_{15:0}$ Δ7 | 1.3 | - | - | - |
| $C_{14:1}$ Δ5 | 0.2 | - | - | - |
| $C_{15:0}$ | 5.5 | 1.5 | 3.8 | 0.9 |
| $C_{15:0}$ DMA | 1.5 | - | - | - |
| $C_{15:0}$ ALDE | 0.1 | - | - | - |
| Anteiso $C_{15:0}$ | 0.2 | - | - | - |
| $C_{15:1}$ Δ7 | - | 7.3 | 11.6 | 1.3 |
| Iso-$C_{15:0}$ | 11.4 | 0.8 | 0.7 | 4.3 |
| Iso-$C_{15:0}$ DMA | 0.4 | - | - | - |
| Iso-$C_{15:1}$ Δ5 | 0.7 | - | - | - |
| Iso-$C_{15:1}$ Δ5+7 | 2.3 | - | - | - |
| Iso-$C_{15:1}$ Δ7 | 2.4 | - | - | - |
| Iso-$C_{15:1}$ Δ9 | 0.3 | - | - | - |
| Iso-$C_{15:1}$ Δ7+9 | - | - | - | 0.9 |
| $C_{16:0}$ | 4.7 | 7.1 | 7.2 | 4.0 |
| Iso-$C_{16:0}$ | - | - | - | 0.7 |
| $C_{16:1}$ Δ7 | 5.2 | 27.5 | 20.6 | 7.0 |
| $C_{16:1}$ Δ7 DMA | 0.7 | - | - | - |
| $C_{16:1}$ Δ? DMA | 0.5 | - | - | - |
| $C_{16:1}$ Δ9 | 3.6 | 4.0 | 2.2 | 1.9 |
| $C_{16:1}$ Δ11 | - | 0.5 | 0.6 | 0.6 |
| Iso-$C_{16:1}$ Δ7 | - | - | 0.4 | 0.4 |
| $C_{17:0}$ | 0.3 | 0.7 | 1.7 | 0.5 |
| $C_{17:0}$ DMA | 0.3 | - | - | - |
| $C_{17:0}$ cyclo Δ9/$C_{17:1}$ Δ? DMA | 5.9 | - | - | - |
| Anteiso-$C_{17:0}$ | - | - | - | 3.1 |
| $C_{17:1}$ Δ7 | - | 2.7 | 6.1 | 2.0 |
| $C_{17:1}$ Δ7/$C_{17:1}$ Δ9/$C_{16:0}$ DMA | 2.6 | - | - | - |
| $C_{17:1}$ Δ9 | - | 10.2 | 12.6 | 1.4 |
| $C_{17:1}$ Δ7+9 | 2.4 | - | - | - |
| $C_{17:1}$ Δ11 | - | 1.1 | 1.5 | - |
| $C_{17:1}$ Δ? DMA | 1.5 | - | - | - |

| | | | | |
|---|---|---|---|---|
| $C_{17:1}$ Δ? DMA | 1.8 | - | - | - |
| Iso-$C_{17:1}$ | 1.8 | - | - | - |
| Iso-$C_{17:1}$ Δ7 | - | 2.0 | 1.7 | 22.3 |
| Iso-$C_{17:1}$ Δ9 | - | 1.0 | 0.6 | 5.6 |
| Iso-$C_{17:1}$ Δ7+9 | 8.5 | - | - | - |
| $C_{18:0}$ | 0.3 | 0.7 | 0.6 | 1.2 |
| $C_{18:1}$ Δ9 | 0.3 | 5.5 | 3.4 | 2.1 |
| $C_{18:1}$ Δ11 | 0.8 | 2.5 | 1.0 | 0.5 |
| Iso-$C_{19:1}$ Δ9 | - | - | - | 0.7 |

* data from [12]
DMA – dimethyl acetal
ALDE - aldehyde
Δ? – position of double-bond unclear
